# ESCRT pathway-dependent MVBs contribute to the morphogenesis of the fungus *Arthrobotrys oligospora*

**DOI:** 10.1101/2022.12.08.519704

**Authors:** Mengqing Tian, Jinrong Huang, Peijie Cui, Guohong Li, Keqin Zhang, Xin Wang

**Author notes:** (GL); (KZ); (XW).

## Abstract

Ammonia opens trap formation in the nematode-trapping (NT) fungus *Arthrobotrys oligospora*, an intriguing morphological switch in NT fungi, where saprophytic mycelia are converted to pathogenic organs. Endocytosis plays a prominent role in nutrient uptake, signaling cascades, and maintenance of cellular homeostasis in higher eukaryotes. Here, we demonstrate that ammonia efficiently promotes endocytosis via the formation of 3D-adhesive mycelial nets in *A. oligospora*. Trap production is followed by the presence of massive multivesicular bodies (MVBs) and membrane rupture and repair. Additionally, both the ubiquitin-proteasome system and the endosomal sorting complex for transport (ESCRT) pathway are immediately linked to endocytosis regulation and MVB formation in ammonia-induced trap formation. Moreover, disruption of the ESCRT-1 complex subunit proteins AoHse and AoVps27 led to the complete loss of membrane endocytosis and trap formation. Finally, the deletion of the deubiquitinase AoSst2 caused a significant reduction in the number of trap structures produced in response to exposure to ammonia or nematodes. Overall, our results increase our knowledge of the molecular mechanisms underlying the phenotypic changes in the NT fungal group, demonstrating that the endocytosis-ESCRT-MVB pathway participates in the regulation of trapping organs.

**Author Summary:** The lifestyle switch of nematode-trapping (NT) fungi is a significant event that increases their pathogenicity to nematode prey, which has resulted in large losses to agricultural crops worldwide. Here, we describe the molecular mechanism underlying how this fungal group forms a NT structure in response to ammonia, a widely preferred nitrogen source in soil niches. Ammonia enhances the endocytosis process, ubiquitin-proteasome system, and endosomal sorting complex for transport (ESCRT) pathway of the model NT fungus *A. oligospora*, thereby generating enriched multivesicular bodies (MVBs) during trap formation. In this process, the cell membrane morphology is remarkably damaged and then repaired. We further found that disruption of the ESCRT-0 subcomplex or ubiquitinase severely blocked trap production and membrane reorganization. Our study provides a new understanding of endocytosis-ESCRT-MVB flux in the transition of fungal NT organs.

## Introduction

Fungi commonly modify their growth style as a survival strategy in response to environmental fluctuations or limited nutrient availability. Pathogenic yeast to filamentous fungi undergoes significant morphological alterations to change from a saprophytic to a carnivorous form during a successful infection of their hosts. Yeast have among the most well-studied pathogenic morphologies, as they can convert between different morphological phenotypes (hyphae, pseudohyphae, and yeast) in response to specific host niches [1, 2]. Similarly, many saprophytic filamentous fungi can sense and respond to their animal or plant hosts to develop infection structures, such as appressoria, haustoria, and traps, depending on which fungi is infecting, parasitizing, or coexisting with their hosts [3-5]. An increasing number of studies have linked fungal morphogenesis to multiple nutrient-response pathways. Limiting ammonium and amino acids can trigger yeast and filamentous fungi to convert to their pathogenic forms through the successful development of pseudohyphae and infection devices [6]. Nematode-trapping (NT) fungi usually grow as saprophytes in ubiquitous soil habitats, trapping nematodes to acquire and survive under starvation conditions. Aside from live nematodes and their glycosides, nitrogen-containing substances such as ammonia or urea also play a significant role in stimulating NT structures to capture and digest nematodes.

Morphological conversion occurs due to extensive remodeling of plasma membranes and cell walls, as well as reorganized cellular composition, producing completely distinct cellular forms [7]. Evidence indicates that many biological processes, from the polarized growth of hyphae into complex reproduction or infection structures, are associated with endocytosis-dependent signal internalization [8]. Regulation of endocytosis plays an instrumental role in nutrient acquisition, signal transduction, and tolerating excess external stimuli [5, 9]. By generating a continuous flow of secretory vesicles, the plasma membrane (PM) receptor sequesters extracellular cues for early endosomes (EE), which fuse with multivesicular bodies (MVB) to be degraded by lysosomes or vacuoles [10, 11]. The endosomal sorting complex required for transport (ESCRT) is required for ubiquitin-dependent degradation of membrane proteins [12]. It is evolutionarily highly conserved across the eukaryotic lineage and is composed of five distinct multi-subunit complexes (ESCRT-0, -I, -II, -III, and the VPS4 complex) that are sequentially recruited to participate in the formation of intraluminal vesicles (ILVs) within MVBs [13]. For an appropriate response to environment cues, ESCRT-0, -I and -II bind to ubiquitinated membrane proteins, whereas ESCRT-III and Vps4 cause ILVs to bud into the endosomal lumen [13, 14]. Overall, MVB sorting machinery is critical for maintaining the correct protein composition and terminating protein function within the lysosome or vacuole [15]. Studies of yeast and mammals have demonstrated that the loss of any ESCRT protein blocks endocytosis-dependent receptor degradation and promotes class E compartment formation, diseases associated with cancer, and neurodegeneration [16, 17]. For the phytopathogenic fungus *Fusarium graminearum*, ESCRT complex are directed toward endocytosis-dependent growth, reproduction, deoxynivalenol production, and full virulence [18]. Moreover, the ESCRT system is involved in hyphal induction and polarity maintenance in *Candida albicans, Ustilago maydis* and other yeast pathogens [19, 20]. Therefore, ESCRT-regulated MVB biogenesis is required for fungal pathogenicity and morphogenesis. The predatory fungi-nematode relationship is one of constitutive coexistence in the soil ecosystem. The fungal species that encounter nematodes as food start forming a variety of infection organs, including adhesive nets/columns/knobs and constricting or non-constricting rings that they use to digest soil-dwelling nematodes to survive and reproduce. Using this interspecies symbiotic model, NT fungi appear to be a potential avenue for the control of plant-parasitic nematodes, which are among the most devastating pests in agriculture [21]. NT structure biogenesis requires a complicated regulatory network in which microRNAs, G-protein signaling, and autophagy are enrolled [22-24]. According to recent research, MVBs can take on an autophagic role, creating a cytoplasmic amphisome that degrades proteins and organelles [25]. In our earlier work, urea-metabolized ammonia was shown to regulate the formation of fungal trap structures in *A. oligospora*, which depend on a specific 3D adhesive network to capture and kill nematodes [26]. Additionally, an ammonia-induced endocytic response has been observed in the elucidation of arrestin family functions in the regulation of trap production in this model NT fungus [27]. Accordingly, many mammalian cells are affected by ammonia during endocytosis in key neurodegenerative diseases [28, 29]. However, the intercellular responses associated with endocytosis and MVB biogenesis in ammonia-led NT traps remain unclear.

To explore the effect of endocytosis on the morphogenesis of NT fungi, we sought to regulate the ESCRT-0 sub-complex (AoVps27/AoHse) and MVB deubiquitinases (DUBs; AoSst2 and AoDoa4) to investigate their effect on intracellular MVB biogenesis, fungal infection devices, and conidiation and growth. We found that ammonia-endocytosis results in the over-accumulation of MVB organelles, allowing *A. oligospora* to produce many trap structures. During the formation of the trapping structures, the fungal cell membrane is subject to hyperactive damage and remodeling.

## Results

### Vesicular trafficking-related pathways exhibit obvious difference according to ammonia levels

Ammonia (NH_3_) is identified as an important signal for the lifestyle transition in NT fungus *A. oligospora* [26]. To explore trap formation by ammonia, we sampled vegetative and ammonia-treated predatory mycelia for transcriptome sequencing. During trap formation, the addition of 25 μM ammonia induced *A. oligospora* mycelia to produce massive 3D adhesive nets (Fig 1A). Transcriptional profiles indicated that pathways related to the whole vesicle system were significantly upregulated during trap production, including ubiquitin-mediated proteolysis, endocytosis, phosphatidylinositol signaling, and SNARE interactions required for vesicular transport (Fig 1B). The endocytosis pathway was significantly upregulated in all periods except for the last stage, in which traps tended to mature (Fig S1). Given that the fluorescent dye FM4-64 is an endocytosis and vesicle trafficking marker in living eukaryotic cells [30, 31], we used this dye to monitor and characterize intracellular response caused by ammonia. When incubated with an ammonia solution, the dye was rapidly absorbed and clustered within intracellular vesicle-like region (Fig 1C, D). Furthermore, because TORC1 is thought to be a regulator of ubiquitin-mediated endocytosis [32, 33], we hypothesized that rapamycin (target of TORC1 kinase) could block the effects of ammonia on trap formation and endocytosis. Our examination confirmed that rapamycin addition suppressed trap production and endocytosis responsiveness after exposure to ammonia (Fig 1E–G). Based on the above data, we believe that ammonia provokes a vesicular trafficking network during trap formation in *A. oligospora*.

**Fig 1.**
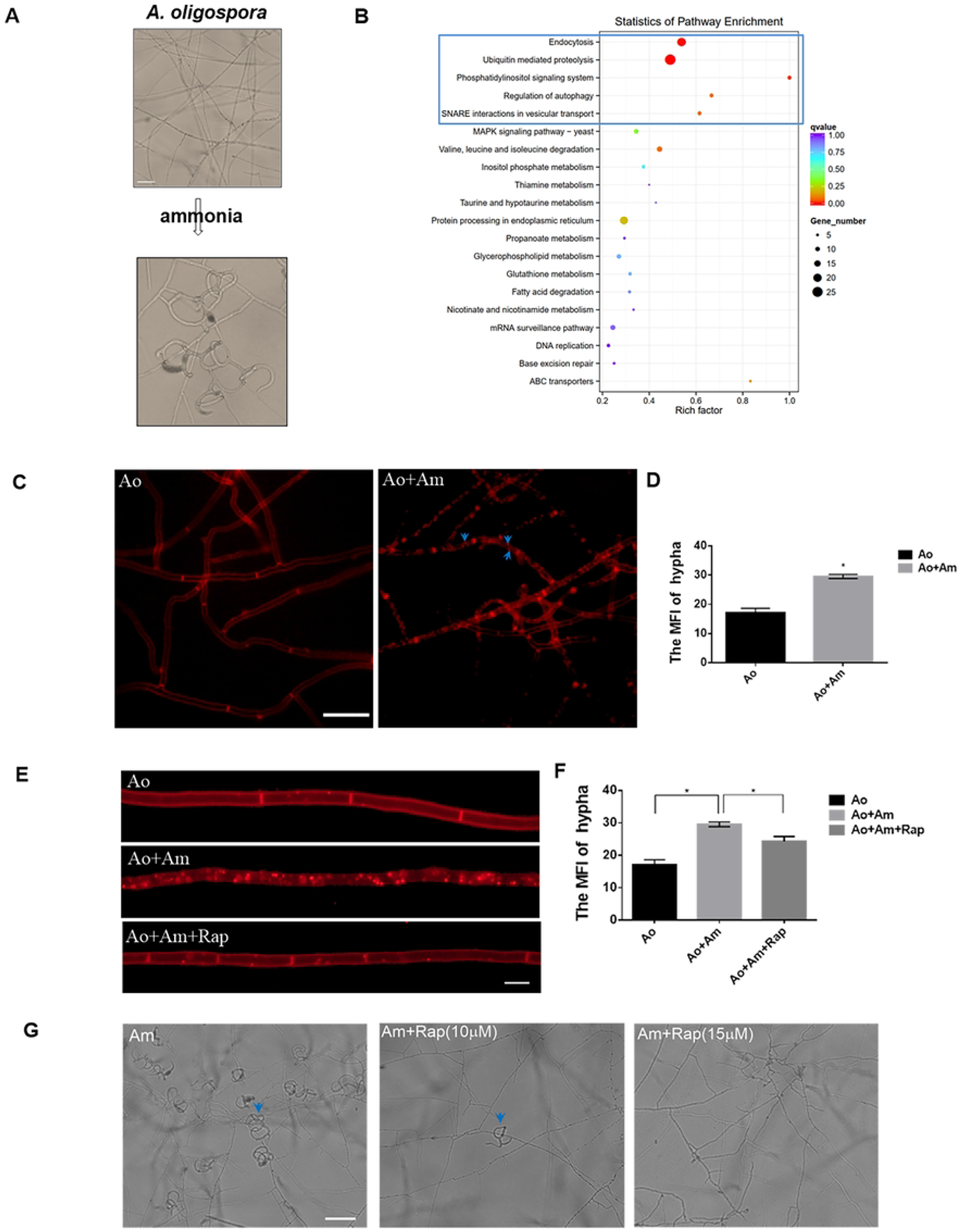
Trap formation and endocytosis in *A. oligospora*. **A**. Traps formation induced by ammonia in *A. oligospora*. Scale bar: 100 µm **B**. KEGG pathway enrichments of up-regulated DEGs induced by ammonia for 4 h, the up-regulated vesicle-related pathways are indicated in the blue box. **C**. Displays of mycelia stained by FM4-64 in control (Ao) and treatment with ammonia (Ao+Am), the blue arrow indicates the puncta formed by dye internalization. Scale bar: 25 µm. **D**. Fluorescence intensity of mycelia stained by FM4-64 in control (Ao) and treatment with ammonia (Ao+Am). **E**. Displays of mycelia stained by FM4-64 in control (Ao) and treatments with ammonia (Ao+Am) and rapamycin (Ao+Am+Rap). Scale bar: 10 µm. **F**. Fluorescence intensity of mycelia stained by FM4-64 in control (Ao) and treatment with ammonia (Ao+Am) and rapamycin (Ao+Am+Rap). **G**. Mycelia treated with ammonia and rapamycin. Scale bar: 200 µm. * P<=0.05(Student’s t test).

### Trap formation is accompanied by MVB dynamics and PM reorganization

Next, to determine whether ammonia affects endosomal phenotype, we analyzed transmission electron microscope (TEM) images of the intracellular endosomes. In general, vegetative hyphal cells have some typical MVB organelles, which contain numerous internalized intraluminal vesicles (ILVs) and they are delivered into the vacuole for further degradation (Fig 2A). However, the ammonia-exposed hyphae accumulated more MVB-like vesicles, which contained fewer ILVs. Remarkably, largely discontinuous membranes were present at the base of the cell wall after 12 h of ammonia treatment (Fig 2B). Intriguingly, once the adhesive net was formed, the damaged membrane was repaired and excessive MVBs disappeared. In addition, many MVB-secreted ILVs were gathered in the PM region, which appeared to be involved in sealing the membrane fissure (Fig 2C). We also observed the surface of the hypha through scanning electron microscope (SEM), which revealed obvious dynamic variation at 0, 12 and 36 h after exposure to ammonia. We observed that the hyphal surface was intact at 0 h (Fig 2D), damaged and broken mycelium fibers appeared at 12 h (Fig 2E), and the hypha was remarkably restored to a complete state at 36 h (Fig 2F). These observations suggested that *A. oligospora* can initiate hyperactive PM reorganization in response to trap development.

**Fig 2.**
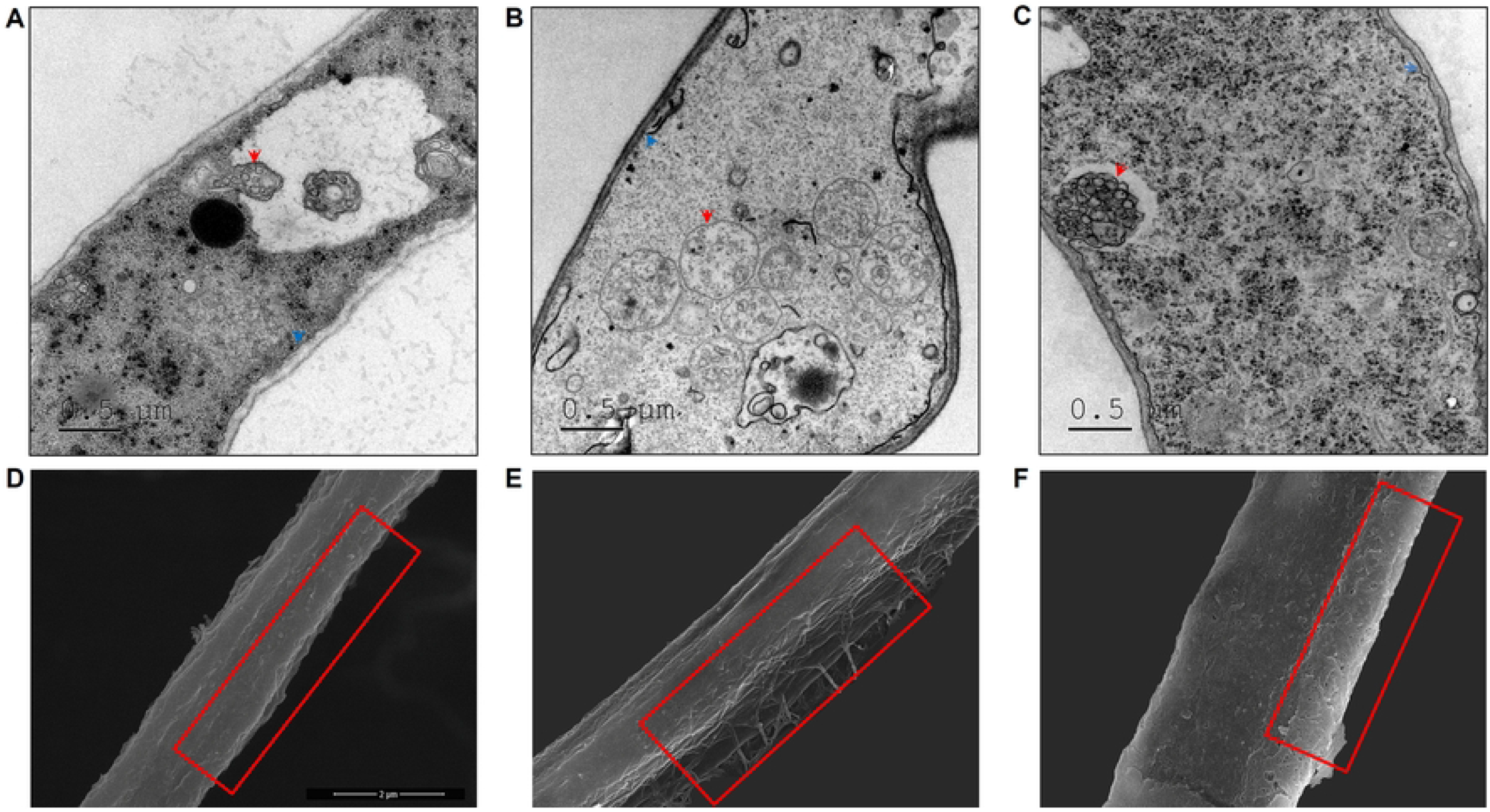
Trap formation is accompanied by MVB dynamics and PM reorganization. **A**. TEM images of intracellular MVB morphology of *A. oligospora* control strain, the red arrow represents MVB morphology in the control cell. Scale bar: 0.5 µm. **B**. TEM images of *A. oligospora* induced by ammonia for 12 h, the red arrow represents MVB morphology in ammonia-treated fungal cell. Scale bar: 0.5 µm. **C**. TEM images of *A. oligospora* induced by ammonia for 36 h, the red arrow represents normal MVB morphology in trap-formed fungal cell. Scale bar: 0.5 µm. **D**. SEM images of the cell wall phenotype of the control *A. oligospora*. Scale bar: 2 µm. **E**. SEM images of the cell wall phenotype of ammonia-treated *A. oligospora* at 12 h, the red box indicates the defective cell wall in the ammonia-treated cell. **F**. SEM images of the cell wall phenotype of ammonia-treated *A. oligospora* at 36 h, the red box indicates the repaired cell wall in the trap-formed cell.

### ECSRT complex and deubiquitinases mediate ammonia-induced trap formation

MVB formation is key to the ubiquitin-mediated proteolysis system (UPS). During this process, the ESCRT pathway sequentially coordinates its diverse subunit proteins to recruit ubiquitin-labeled proteins into the MVB, thus maintaining protein homeostasis [34, 35]. The ESCRT complex is highly conserved between yeast and filamentous fungi and comprises four complexes (ESCRT-0, -I, -II, -III, and VPS4) containing approximately 30 proteins [36]. Based on transcriptional profile, ammonia differentially upregulated the expression of the ECSRT complex and DUBs (Fig 3A; Table S1). RT-PCR assay also showed that the ESCRT-coding genes (*AoVps27, AoHse, AoVps23, AoVps36, AoVps4, AoSnf7*) and DUB-coding genes (*AoSst2* and *AoDoa4*) were significantly upregulated by over 6- to 20-fold after the addition of ammonia (Fig 3B). Because deubiquitination acts in concert with ubiquitination in organisms, we tested the interaction of DUBs (AoSst2 and AoDoa4) with the ESCRT-0 subunit (AoHse/AoVps27). The yeast two-hybrid (Y2H) assay revealed that AoVps27 couples with AoHse, as well as with the DUBs AoSst2 and AoDoa4 (Fig 3C). Furthermore, concerted expression was observed in *A. oligospora* mutants of Δ*AoVps27* and Δ*AoHse*, and the destruction of one gene reduced the expression of the other or DUB-related genes, that is the expression levels of *AoSst2* and *AoDoa4* were significantly downregulated in mutants Δ*AoHse* and Δ*AoVps27* (Fig 3D, E). These analyses suggest that ESCRT-0 and DUBs may work together in trap formation via ammonia signaling.

**Fig 3.**
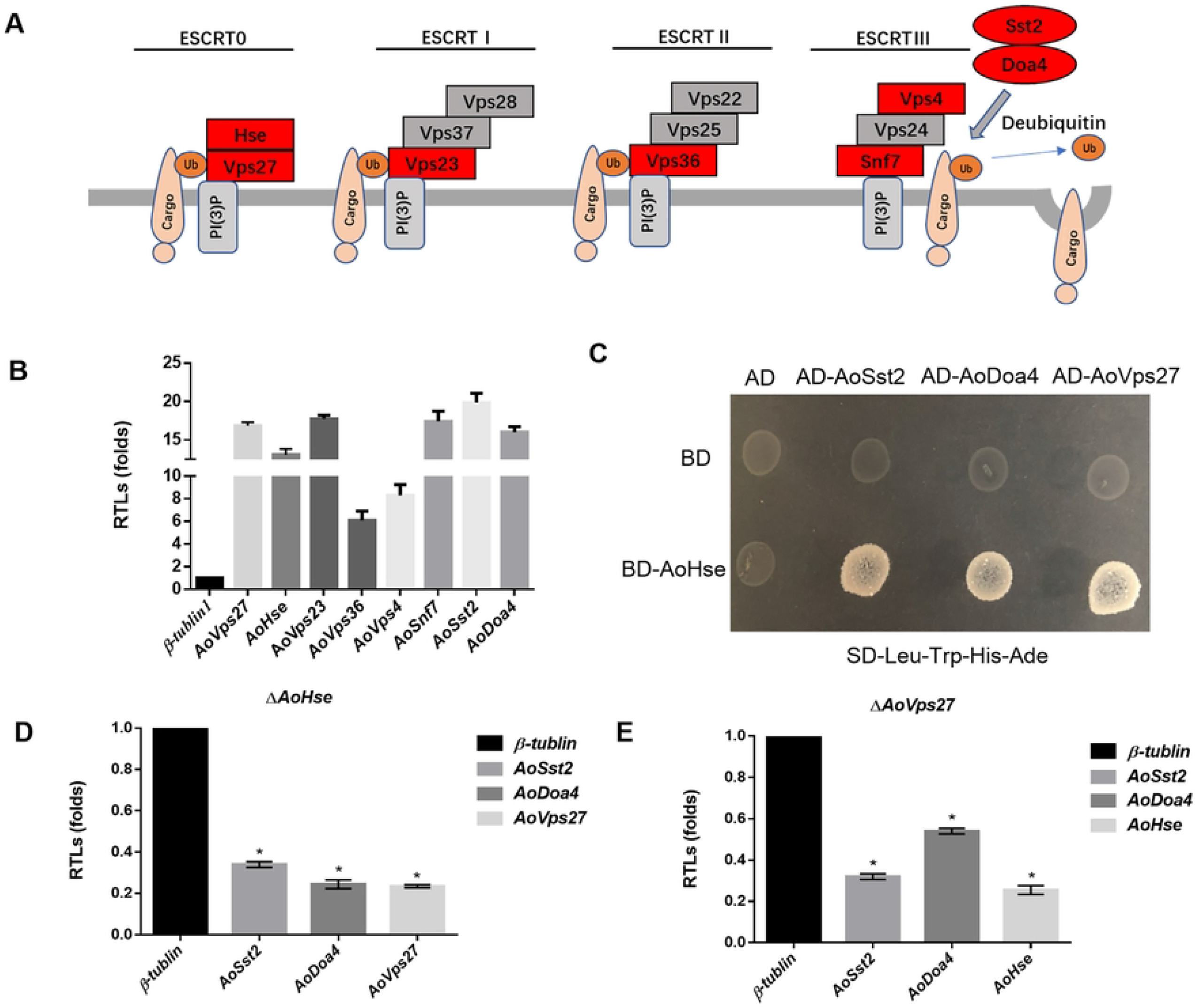
Relations between ECSRT-0 and deubiquitinases. **A**. Structure schematics of ESCRT complex. **B**. Transcriptional levels of genes involved in ESCRT-0 and deubiquitinases when exposure to ammonia. **C**. Y2H results about ECSRT-0 and deubiquitinases. **D**. Transcriptional levels of genes *AoVps27, AoSst2* and *AoDoa4* in mutant Δ*AoHse*. **E**. Transcriptional levels of genes *AoHse, AoSst2* and *AoDoa4* in mutant Δ*AoVps27*. * P<=0.05(Student’s t test).

### Trap formation is linked to ESCRT-0-mediated MVB biogenesis and PM repair

ESCRT-0 is the most upstream sub-complex of the ESCRT pathway and is responsible for capturing ubiquitinated proteins in the MVB-lysosome/vacuole degradation pathway. Evolutionary tree analysis showed that the ESCRT-0 complex (Hse-Vps27) is highly conserved from *Saccharomyces cerevisiae* to human. The two ESCRT-0 proteins contain more than one ubiquitin interaction motif (UIM) that binds to ubiquitinated molecules (Fig S2). We knocked out the genes to examine the effects of AoHse and AoVps27 on trap formation. Fragments of *AoHse* and *AoVps27*, 300 bp and 753 bp in length, respectively, were replaced by hygromycin-encoding genes; none of them were subsequently detected in the corresponding mutants (Fig S3). In the presence of ammonia or nematodes, both of the mutation completely blocked trap formation (Fig 4A, B). Consequently, the fungus is no longer able to capture and kill nematode preys (Fig S4). Additionally, we examined FM4-64 staining and TEM images to determine whether the loss of traps was related to changes in endocytosis or MVB traffic. There was a clear inhibition of FM4-64 uptake in mutant hyphal cells after 10 min of ammonia exposure; further, mutant mycelium fluorescence intensity (MFI) was significantly reduced compared to WT mycelium (Fig 4C, D).

**Fig 4.**
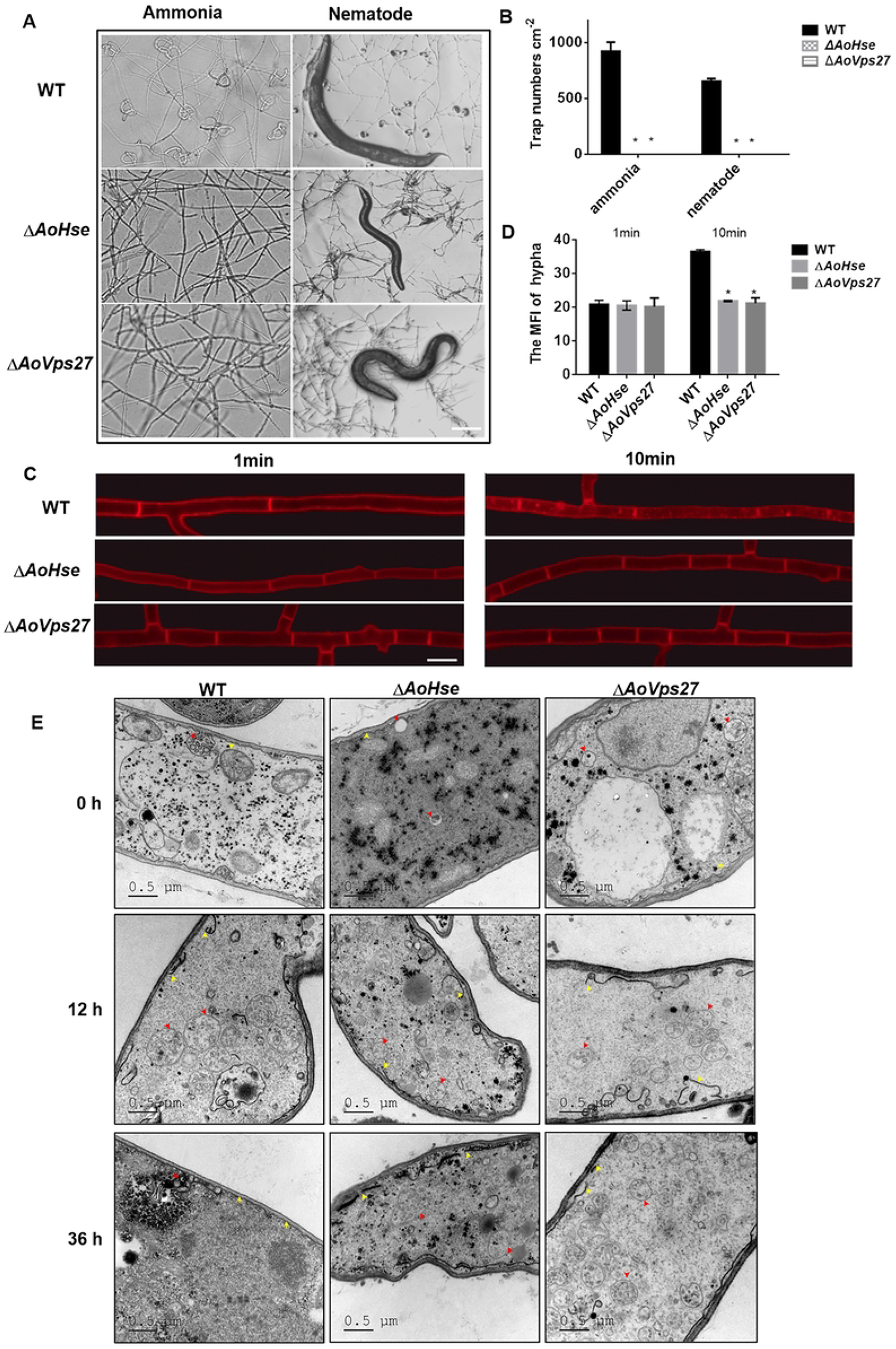
Related phenotypic in mutants Δ*AoHse* and Δ*AoVps27*. **A**. Traps formation induced by ammonia in WT and mutants Δ*AoHse* and Δ*AoVps27*. Scale bar: 200 µm. **B**. Traps numbers in WT and mutants Δ*AoHse* and Δ*AoVps27* when exposure to ammonia and nematodes. **C**. Displays of mycelia stained by FM4-64 in WT and mutants Δ*AoHse* and Δ*AoVps27*. Scale bar: 25 µm. **D**. Fluorescence intensity of mycelia stained by FM4-64 in WT and mutants Δ*AoHse* and Δ*AoVps27*. **E**. TEM images of WT and mutants Δ*AoHse* and Δ*AoVps27* when exposure to ammonia for 0, 12 and 36 h (Red arrow refers to MVBs and yellow arrows refers to membranes). Scale bar: 0.5 µm. * P<=0.05(Student’s t test).

To further determine whether MVBs co-occur with cell membrane integrity, we compared vesicle morphology and membrane morphology in WT and mutant strains. Based on TEM observation, the mutants showed that the mutants had severe MVB defect, where the MVBs are absent in favor of a few small empty vesicles (Fig 4E). Besides, In the Δ*AoHse* mutant, numerous small vesicles had accumulated in the vacuole-like region, while the Δ*Aovps27* vacuole appeared to be smaller than the WT vacuole under ammonia treatment (Fig S5). The above description suggested that an aberrant MVB-vacuolar pathway may contribute to the loss of trap in ESCRT-0. As shown in Fig 2C, the PM morphology was intact when the trap was formed following 36-h treatment with ammonia. Thus, we investigated whether ESCRT-0 is also involved in membrane remodeling. Similar to WT, mutant strains Δ *AoHse* and Δ *AoVps27* showed severe damage to their embranes after 12 h of ammonia exposure; however, after 36 h, membrane disruption persisted (Fig 4E). In light of the above, ESCRT-0 mutations disrupt PM remodeling and MVB biogenesis in response to ammonia signals.

### Deubiquitination is key for the fitness of MVB in *A. oligospora*

Given the previously mentioned conserved interaction between ESCRT-0 and DUBs, we hypothesized that blocking DUBs AoSst2 and AoDoa4 would disrupt MVB biogenesis and reduce trap formation in *A. oligospora*. Based on the corresponding protein sequences analysis, AoSst2 and AoDoa4 exhibit a high degree of conservation (Fig S6). We knocked out the genes to examine the effects of *AoSst2* and *AoDoa4* on trap formation. Length of 572 bp and 813 bp of fragments of *AoSst2* and *AoDoa4* respectively were replaced by hygromycin encoding genes in genes, neither were detectable in the corresponding mutant (Fig S7). As expected, the absence of the DUBs AoSst2 and AoDoa4 inhibited trap production. This was particularly pronounced for Δ*AoSst2*, which almost failed to form traps from ammonia induction (Fig 5A). Respectively, the number of traps decreased by 74.8% and 34.1% at 48 h after ammonia induction and decreased 54.8% and 14.1% at 48 h after nematode induction (Fig 5 B, Fig S8B). Meanwhile, nematode capture efficiency of Δ*AoSst2* and Δ*AoDoa4* slightly decreased by 22.3% and 4.1%, respectively (Fig S8A). Furthermore, we compared endocytosis changes between the strains and found that the deleting of *AoSst2* caused obvious delays in endocytosis, but that the *AoDoa4* mutation did not result in similar defects (Fig 5 C, D). Based on our observation of the MVB phenotype, we found that DUB deletion also caused some empty and small vesicle fragments to occur inside the cell (Fig 5E). These data suggest that deubiquitination regulates trap formation in *A. oligospora*.

**Fig 5.**
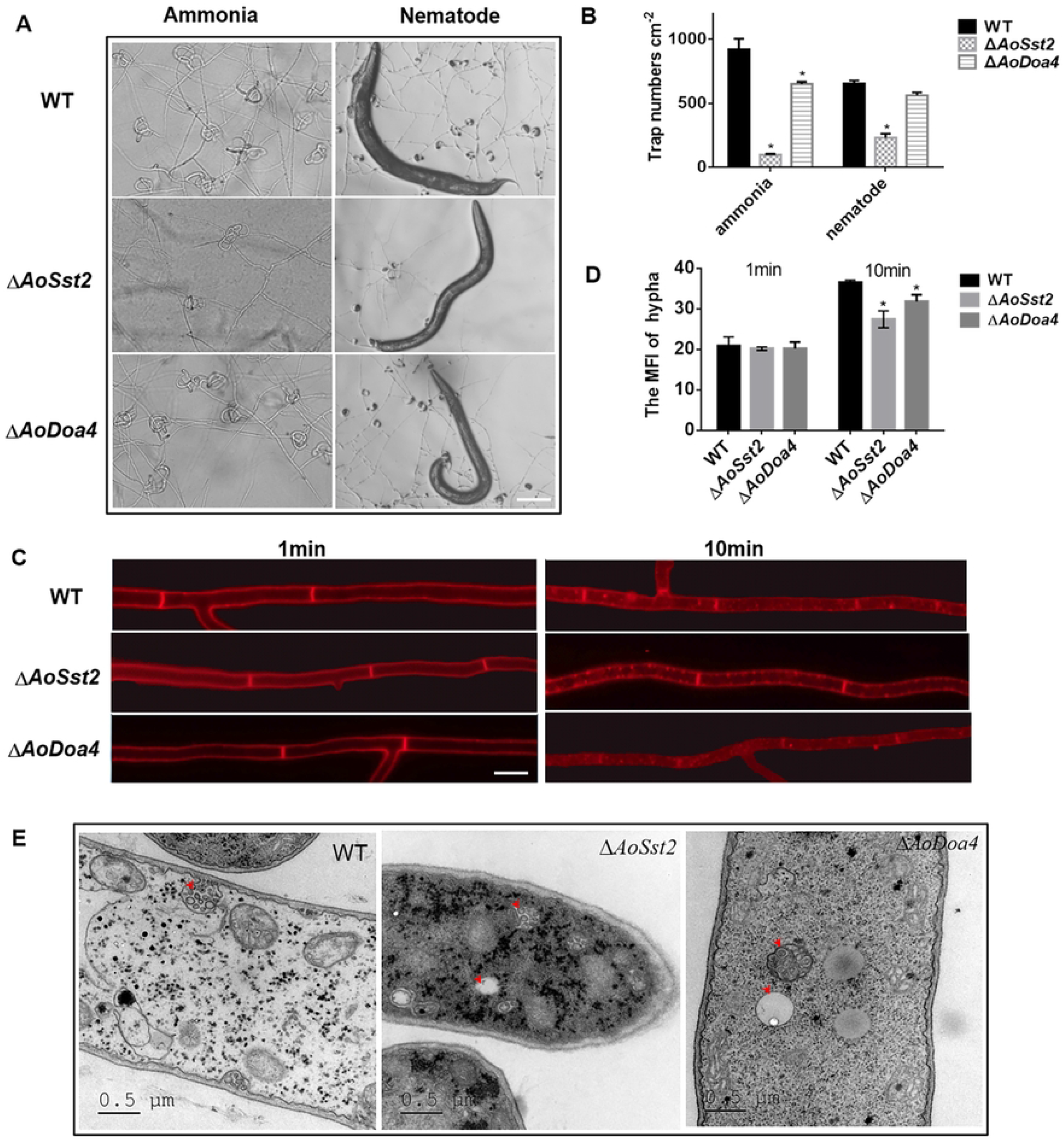
Related phenotypic in mutants Δ *AoSst2* and Δ *AoDoa4*. **A.** Traps formation induced by ammonia in mutants Δ *AoSst2* and Δ *AoDoa4*. Scale bar: 200 µm. **B**. Traps numbers in WT and mutants Δ *AoSst2* and Δ *AoDoa4* when exposure to ammonia and nematodes. **C**. Displays of mycelia stained by FM4-64 in mutants Δ *AoSst2* and Δ *AoDoa4*. Scale bar: 25 µm. **D**. Fluorescence intensity of mycelia stained by FM4-64 in mutants Δ*AoSst2* and Δ*AoDoa4*. **E**. TEM images of WT and mutants Δ*AoHse* and Δ*AoVps27* when exposure to ammonia for 0h, 12h and 36h (Red arrow refers to MVBs). Scale bar: 0.5 µm * P<=0.05(Student’s t test).

## Discussion

Fungal traps differentiate constitutively in response to nematodes [37]. Diverse nematodes always coexist with predatory fungi, which tend to eavesdrop on nematode-produced cues to build infection weapons to catch and digest their nematode preys[38, 39]. NT fungi predate nematodes for nutrients, particularly nitrogen sources [23]. In addition to the nematode itself and its metabolite ascarosides, the nitrogenous compounds (amino acids, ammonia and urea) can also stimulate a change in fungal lifestyle from saprophytic to parasitic[40]. The investigation of trap regulation will lead to comprehensive understandings of how cell types transform and why NT fungi have co-evolved with the emergence of nematodes. Over the past decade, G-protein-associated signaling has been considered the main explanation underlying the trap formation [23, 41, 42]. Multi-omics analysis commonly revealed that the membrane biogenesis-associated pathways contribute to trap formation [43]. Here, we discovered that endocytosis and vesicle routines are involved in signal ammonia-triggered trap formation in the NT fungus *A. oligospora* (Fig 6).

**Fig 6.**
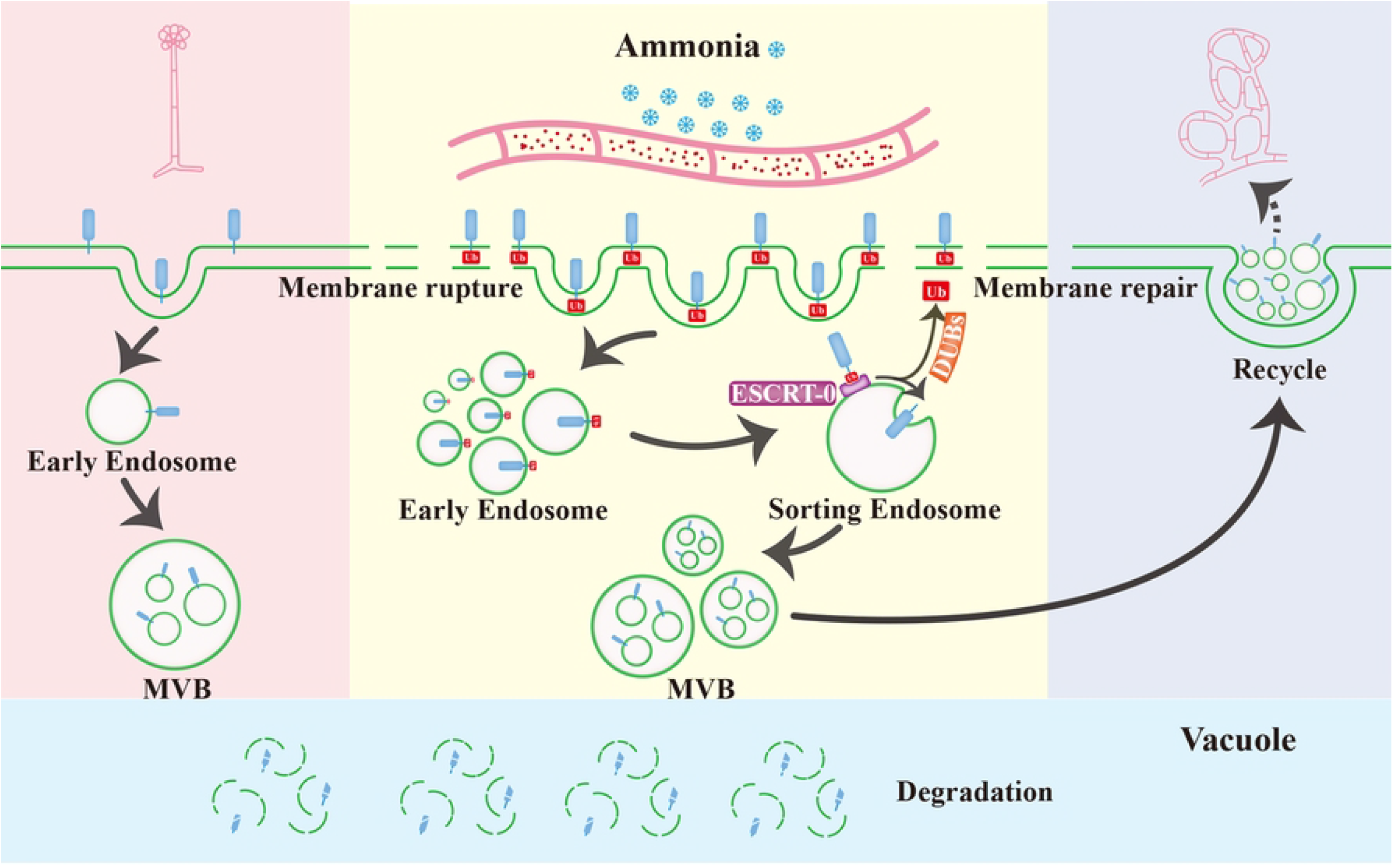
Proposed model for the endocytosis-ESCRT-MVB contributes to morphogenesis of the fungus *Arthrobotrys oligospora*. Ammonia stimulation first triggers higher PM endocytosis, thereby generating more MVBs regulated by the ESCRT-0 complex (AoHse/AoVPS27) and the interactors Dubiquitinases AoSst2, and AoDoa4. Increased endocytosis may result in discontinuous membrane organization. Additional to being destined for vacuolar degradation, the excess MVBs are also recycled to the PM region to help reseal the PM wounds. Eventually, the fungus produces 3D-adhesive nets under the regulation of endocytosis-ESCRT-MVB flux induced by ammonia.

First, PM endocytosis mainly occurs during signal transduction and reconstruction of cell polarity; however, ammonia-induced traps are also produced through PM endocytosis. Endocytosis is a cellular response to a multitude of stress conditions that allows organisms to tolerate complicated conditions by shutting down external signaling [44]. Similar physiological changes have been widely identified in mammals and fungal cells such as *Aspergillus nidulans* and yeast [45-47]. We also previously described that ammonia induces trap formation by regulating endocytosis through arrestin-GPCR signaling [27].

Second, a sharp accumulation of endosomes within the cell may disrupt the integrity of the PM. However, following the disappearance of excessive endosomes and the repair of plasma membrane, the fungus produces many adhesive trapping structures. These phenomena are similar to the biological event occurred in *Xenotoca eiseni*, where irregularly delineated giant endosomes appear in embryos stressed by dose exceeding 10 mM NH_4_Cl [48]. The increase in the number of endosomes may be due to the highly damaged membrane caused by ammonia treatment, as the primary assembly of EE is determined by the invagination or internalization of PMs. When faced with external chemicals or pathogens, organisms must depend on their cellular membranes integrity to maintain cellular homeostasis [49]. Numerous vesicles were gathered at the PM lesion region, which may be a cell membrane repair strategy to repair the cell membrane[50]. By stimulating cells with ammonia, traps may form more frequently in response to membrane damage, which further help the fungi survive in another growth mode. In this process, the higher ubiquitin-mediated protein degradation pathway leads to more membrane proteins or misfolded proteins being internalized or enclosed within the MVB. Here, we suggest that the degradation triggers a sharp cellular starvation response due to the fluctuation of the endo-vacuolar system. For NT fungi, starvation is the switch signal from vegetative growth to a predatory lifestyle [51].

Third, in light of the substantial role that the ESCRT plays in plasma membrane fission and repair [52], we believe that the differentiation of fungal trap structures is followed by the assembly of the ESCRT and that membrane reorganization may constitute the initiation signal for trap biogenesis. Having an array of small vesicles gathered between the rapture-like region and the cell wall is not just an accident; rather, it is more likely that the vesicles are retained to maintain membrane integrity and set up a trap. This study demonstrates that the loss of function of ESCRT-0 sub-protein completely prevents membrane repair, resulting in a failure of trap production. This is the direct evidence that ESCRT apparatus is responsible for the membrane repair upon ammonia stress. The ESCRT machinery is an ancient, evolutionarily conserved membrane remodeling complex deployed by cells to perform a diverse collection of physiological and pathophysiological processes. ESCRT proteins are required for MVB biogenesis. More recently, ESCRTs have been shown to play essential roles in repairing damaged cellular membranes, thus preserving cellular viability and organellar function [53]. The development of traps allows for the expression of most of ESCRT genes. Each ESCRT-0 protein contained a UIMs array that was responsible for the recognition of UB-bound protein molecules. Additionally, ubiquitination is a reversible post-translational modification in which DUBs hydrolyze Ub-substrate and Ub-Ub isopeptide bonds. Thus, ubiquitinase and DUBs are master regulators within the Ub system that impinge on diverse cellular processes[54]. In our study, we deleted DUBs AoSst2 and AoDoa4 to examine the effect of deubiquitination on trap formation. The current results illustrate that the deubiquitination is involved in endosome formation, membrane repair and trap formation.

In summary, our results suggest details about two main events before trap formation: first, that the upregulation of endocytosis is vital to endosome enrichment, which activates protein degradation, and the fungal cell enters starvation condition. Second, wounds in the membrane may be due to the overuse of membrane components in the endosomal assembly, which cannot be supplemented in time, and the PM integrity-dependent generation of traps by ammonia. These data are consistent with the aSMase-mediated repair mechanism in mammalian cells. This is the first report showing that NT fungal species has the machinery to repair ESCRT-mediated PM damage to promote the morphogenesis of trap organs. Nevertheless, the TEM image highlights an important event: the amounts of vesicle recycled into the periplasmic space clearly exceeds that needed for sealing the membrane wounds. As part of our future study, we will discuss whether and how the exocytosis vesicle mediates trap development in this fungus.

## Materials and Methods

### Fungal Strains and Culture Conditions

Fungus *A. oligospora* ATCC 24927 is stored in the State Key Laboratory for Conservation and Utilization of Bio-Resources in Yunnan and it grows on the standard nutrient medium of PDA (20% potato, 2% dextrose, and 1.5% Agar) medium at 28°C. *S. cerevisiae* FY834 is cultured in YPD (1% yeast, 2% peptone, and 2% dextrose) broth or on the solid YPD medium (YPD supplemented with 1.5% agar), and the recombinant yeast is selected from SC-Ura medium [55]. Y2H Gold yeast strain is cultured in YPDA (TaKaRa Bio,Shiga, Japan). *Escherichia coli* strain DH5α is used to preserve the plasmids pRS426, pCSN44, pGADT7 and pGBKT7. Additionally, liquid TG (1% tryptone, and 1% glucose) medium is suitable for mycelial collection, PDAS medium (PDA supplemented with 10 g/L molasses and 0.4 M saccharose) is used for protoplast regeneration, solid CMY (2% maize boiled, 0.5% yeast extract, 1.5% agar) was used for conidia culture, WA (deionized water with 1.5% agar) is suitable for traps-inducing tests. The nematode *Caenorhabditis elegans* is maintained on NGM medium at 20 °C [56].

### Sample collection and transcriptome sequencing analysis

The NT fungus *A. oligospora* was incubated on CMY medium at 28 °C for 10 days and the conidia were collected. Subsequently, about 10^4^ conidia were spread on the WA medium covered with cellophane in 9cm plate and maintained at 28 °C for 48 h. Then, 25 μM-ammonia was added to each plate for the trap induction at 28 °C. Normally, traps could form within 36 h. In order to obtain the active regulation pathways of trap formation during the induction process, we collected mycelia treated for 4, 12, 36 and 48 h and the control mycelia for transcriptome sequencing analysis (Novogene Co. Ltd).

### Quantitative real-time PCR (RT-PCR) analysis

To verify the gene expression difference in transcriptome sequence data, samples were prepared as before for the real time PCR analysis of some significantly upregulated genes. Besides, WT and the mutants of Δ*AoHse*, Δ*AoVps27*, Δ*AoSst2* and Δ*AoDoa4* were incubated on PDA medium at 28 °C and the hyphae were collected for the determination of transcript levels of genes involved in sporulation and MVBs biogenesis. Total RNA of all samples was extracted using an RNA Extraction Kit (QIAGEN, Germany); then, the corresponding cDNA was transcribed using a Fast Quant RT Kit with gDNA (TaKaRa Bio, Shiga, Japan). The cDNA was used as a template for RT-PCR assays, which were conducted to analyze the transcriptional levels of candidate genes. The gene-specific RT-PCR primers (Table S2) were designed with Primer5 software, and the β-tubulin-encoding gene (Tub, *AOL_s00076g640*) was used as an internal standard. RT-PCR was performed in triplicate for each gene as previous described [57], and fold-changes were calculated using the 2^-ΔΔCt^ method [58].

### Endocytosis analyses

To test endocytosis, the fungus blocks of the WT and mutant strains were cultured on two-layer water agar (WA) plates for 3 days; then, the upper medium was cut and these fungal blocks were stained with FM4-64 (Biotium, California, USA) for 1 min and 10 min, respectively. Then, the samples were washed with ddH_2_O for three times and observed with a fluorescence microscope (Nikon, Tokyo, Japan) [59]. To observe the changes of MVBs and cell members, samples were prepared for TEM and SEM analysis, ultrathin sections of hyphal cells were prepared and examined as previously described [60].

### Yeast two-hybrid analyses

Protein-protein interactions were assayed with the Matchmaker yeast two-hybrid system (Clontech, Mountain View, CA). ORFs of *AoHse* was amplified from first-strand cDNA of *A. oligospora* and cloned into pGBK7 (Clontech) as the bait constructs. For the *AoVps27, AoSst2* and *AoDoa4*, their ORFs were amplified and cloned into pGADT7 as the prey constructs. The resulting bait and prey vectors were co-transformed in pairs into yeast strain Gold yeast (Clontech). The SD-Leu-Trp transformants were isolated and assayed for growth on SD-Leu-Trp-His-Ade medium [61].

### Deletion of the genes *AoHse, AoVps27, AoSst2* and *AoDoa4*

The target genes were disrupted by homologous recombination[55]. Two flanking fragments were amplified by PCR with paired primers 5F/5R, 3F/3R (Table S2) with *A. oligospora* genomic DNA as the template, and the hygromycin B resistance cassette (*hyg* cassette) were amplified using the primer pair hyg-F/hyg-R from the plasmid pCSN44. The three fragments together with a linearized vector pRS426(digested with EcoRI/XhoI) were transform to *S. cerevisiae* strain FY834 cells to get the final recombinant plasmid pRS426-gene-hyg. The deletion cassette was amplified by PCR with paired primers 5F/3R. Then, protoplasts were prepared from the fungus *A. oligospora*, and the purified deletion fragments were transformed into the protoplasts with PEG-mediated approach method. Putative mutant colonies were selected on PDAs medium containing 200 μg/mL hygromycin B and confirmed by PCR amplification using specific primers [62].

### Trap formation and pathogenicity assays

The WT and mutant strains were incubated on CMY medium at 28 °C for 10 days and conidia were collected. Subsequently, 10^4^ conidia of the WT and mutant strains were respectively spread on WA plates and maintained at 28 °C for 48 h. Then, 200-300 nematodes were added to each plate to induce trap formation at 28 °C, and observed and recorded the numbers of traps and captured nematodes at 24, 36 and 48 h. Similarly, when the conidia maintained on WA for 48 h at 28 °C, 25 μM ammonia was added to each plate to induce trap formation at 28 °C, and observed and recorded the number of traps at 36, 48, and 60 h.

## Funding

This work is supported by the National Natural Science Foundation of China (32260029), the Applied Basic Research Foundation of Yunnan Province (202101AT070266) to Xin Wang, the postdoctoral science foundation (2022MD723850) to Mengqing Tian. The funders had no role in study design, data collection and analysis, decision to publish, or preparation of the manuscript.

## Competing Interests

The authors have declared that no competing interests exist.

## Supporting information captions

**S1 Fig** KEGG pathway enrichments of up-regulated DEGs induced by ammonia for 12h, 36h and 48h

**S2 Fig** Evolutionary tree analysis and structure diagram of AoHse and AoVps27 **S3 Fig** Diagrams for the disruptions of genes *AoHse* and *AoVps27* and verification of mutants

**S4 Fig** Nematodes capture ability in mutants Δ*AoHse* and Δ*AoVps27*

**S5 Fig** Vacuoles represented in mutants Δ*AoHse* and Δ*AoVps27*

**S6 Fig** Evolutionary tree analysis and structure diagram of AoSst2 and AoDoa4

**S7 Fig** Diagrams for the disruptions of genes *AoSst2* and *AoDoa4* and verification of mutants

**S8 Fig** Nematodes capture ability in mutants Δ*AoSst2* and Δ*AoDoa4*

**S1 Table** Genes upregulated in endocytosis pathway reduced by ammonia

**S2 Table** List of primers used in this study

**S3 Table** Homologous proteins of AoHse, AoVps27, AoSst2 and AoDoa4

## Notes

### Competing Interest Statement

The authors have declared no competing interest.

## References

1. Li Z., and Nielsen K., Morphology Changes in Human Fungal Pathogens upon Interaction with the Host. Journal of Fungi, 2017. 3(4):66.

2. Kadosh D., Shaping up for battle: morphological control mechanisms in human fungal pathogens. PLoS Pathog, 2013. 9(12):e1003795.

3. Chethana K., Jayawardena R.S., Chen Y.-J., Konta S., Tibpromma S., Abeywickrama P.D. et al., Diversity and function of appressoria. Pathogens, 2021. 10(6):746.

4. Dar S.A., Rather B.A., and Kandoo A.A., Insect pest management by entomopathogenic fungi. J Entomol Zool Stud, 2017. 5:1185–1190.

5. Zhang Y., Li S., Li H., Wang R., Zhang K.Q., and Xu J., Fungi– nematode interactions: diversity, ecology, and biocontrol prospects in agriculture. Journal of Fungi, 2020. 6(4):206.

6. Boeckstaens M., André B., and Marini A.M., The yeast ammonium transport protein Mep2 and its positive regulator, the Npr1 kinase, play an important role in normal and pseudohyphal growth on various nitrogen media through retrieval of excreted ammonium. Molecular microbiology, 2007. 64(2):534–546.

7. Lin X., Alspaugh J.A., Liu H., and Harris S., Fungal morphogenesis. Cold Spring Harbor perspectives in medicine, 2015. 5(2):a019679.

8. Riquelme M., Aguirre J., Bartnicki-García S., Braus G.H., Feldbrügge M., Fleig U. et al., Fungal morphogenesis, from the polarized growth of hyphae to complex reproduction and infection structures. Microbiology and Molecular Biology Reviews, 2018. 82(2):e00068–00017.

9. Apodaca G., Modulation of membrane traffic by mechanical stimuli. American Journal of Physiology-Renal Physiology, 2002. 282(2):F179–F190.

10. Piper R.C., Multivesicular bodies: biogenesis and function. 2013.

11. Singh M.K., Krüger F., Beckmann H., Brumm S., Vermeer J.E., Munnik T. et al., Protein delivery to vacuole requires SAND protein-dependent Rab GTPase conversion for MVB-vacuole fusion. Current Biology, 2014. 24(12):1383–1389.

12. Frankel E., and Audhya A., ESCRT-dependent cargo sorting at multivesicular endosomes. In: Seminars in cell & developmental biology: 2018. Elsevier: 4–10.

13. Schmidt O., and Teis D., The ESCRT machinery. Current Biology, 2012. 22(4):R116–R120.

14. Migliano S.M., and Teis D., ESCRT and membrane protein ubiquitination. Endocytosis and Signaling, 2018.107–135.

15. Davies B.A., Lee J.R., Oestreich A.J., and Katzmann D.J., Membrane protein targeting to the MVB/lysosome. Chemical reviews, 2009. 109(4):1575–1586.

16. Mattissek C., and Teis D., The role of the endosomal sorting complexes required for transport (ESCRT) in tumorigenesis. Molecular membrane biology, 2014. 31(4):111–119.

17. Stuffers S., Brech A., and Stenmark H., ESCRT proteins in physiology and disease. Experimental cell research, 2009. 315(9):1619–1626.

18. Xie Q., Chen A., Zhang Y., Yuan M., Xie W., Zhang C. et al., Component Interaction of ESCRT Complexes Is Essential for Endocytosis-Dependent Growth, Reproduction, DON Production and Full Virulence in Fusarium graminearum. Frontiers in microbiology, 2019. 10:180–180.

19. Yang T., Li W., Li Y., Liu X., and Yang D., The ESCRT System Plays an Important Role in the Germination in Candida albicans by regulating the expression of hyphal-specific genes and the localization of polarity-related proteins. Mycopathologia, 2020. 185(3):439–454.

20. Haag C., Klein T., and Feldbrügge M., ESCRT mutant analysis and imaging of ESCRT components in the model fungus Ustilago maydis. In: The ESCRT Complexes. Springer; 2019: 251–271.

21. Sato K., Kadota Y., and Shirasu K., Plant immune responses to parasitic nematodes. Frontiers in plant science, 2019.1165.

22. Ji X., Li H., Zhang W., Wang J., Liang L., Zou C. et al., The lifestyle transition of Arthrobotrys oligospora is mediated by microRNA-like RNAs. Science China Life Sciences, 2020. 63(4):543–551.

23. Bai N., Zhang G., Wang W., Feng H., Yang X., Zheng Y. et al., Ric8 acts as a regulator of G-protein signalling required for nematode-trapping lifecycle of Arthrobotrys oligospora. Environmental Microbiology, 2022. 24(4):1714–1730.

24. Zhou D., Zhu Y., Bai N., Yang L., Xie M., Yang J. et al., AoATG5 plays pleiotropic roles in vegetative growth, cell nucleus development, conidiation, and virulence in the nematode-trapping fungus Arthrobotrys oligospora. Science China Life Sciences, 2022. 65(2):412–425.

25. Fader C., and Colombo M., Autophagy and multivesicular bodies: two closely related partners. Cell Death & Differentiation, 2009. 16(1):70–78.

26. Wang X., Li G.-H., Zou C.-G., Ji X.-L., Liu T., Zhao P.-J. et al., Bacteria can mobilize nematode-trapping fungi to kill nematodes. Nature communications, 2014. 5(1):1–9.

27. Zhou L., Li M., Cui P., Tian M., Xu Y., Zheng X. et al., Arrestin-coding genes regulate endocytosis, sporulation, pathogenicity, and stress resistance in Arthrobotrys oligospora. Frontiers in cellular and infection microbiology, 2022. 12.

28. Seiler N., Ammonia and Alzheimer’s disease. Neurochemistry international, 2002. 41(2-3):189–207.

29. Hirano S., and Kanno S., Macrophage receptor with collagenous structure (MARCO) is processed by either macropinocytosis or endocytosis-autophagy pathway. PloS one, 2015. 10(11):e0142062.

30. Steinberg G., Endocytosis and early endosome motility in filamentous fungi. Current opinion in microbiology, 2014. 20:10–18.

31. Van Gisbergen P., Esseling-Ozdoba A., and Vos J., Microinjecting FM4–64 validates it as a marker of the endocytic pathway in plants. Journal of Microscopy, 2008. 231(2):284–290.

32. MacGurn J.A., Hsu P.C., Smolka M.B., and Emr S.D., TORC1 regulates endocytosis via Npr1-mediated phosphoinhibition of a ubiquitin ligase adaptor. Cell, 2011. 147(5):1104–1117.

33. Boeckstaens M., Llinares E., Van Vooren P., and Marini A.M., The TORC1 effector kinase Npr1 fine tunes the inherent activity of the Mep2 ammonium transport protein. Nature communications, 2014. 5(1):1–12.

34. Davies C.W., Paul L.N., and Das C., Mechanism of recruitment and activation of the endosome-associated deubiquitinase AMSH. Biochemistry, 2013. 52(44):7818–7829.

35. Zhu W., Zheng D., Wang D., Yang L., Zhao C., and Huang X., Emerging roles of ubiquitin-specific protease 25 in diseases. Frontiers in Cell and Developmental Biology, 2021. 9:698751.

36. Juan T., and Fürthauer M., Biogenesis and function of ESCRT-dependent extracellular vesicles. In: Seminars in cell & developmental biology: 2018. Elsevier: 66–77.

37. Yang C.T., Vidal-Diez de Ulzurrun G., Gonçalves A.P., Lin H.C., Chang C.W., Huang T.Y. et al., Natural diversity in the predatory behavior facilitates the establishment of a robust model strain for nematode-trapping fungi. Proceedings of the National Academy of Sciences, 2020. 117(12):6762–6770.

38. Chen S.A., Lin H.C., Schroeder F.C., and Hsueh Y.P., Prey sensing and response in a nematode-trapping fungus is governed by the MAPK pheromone response pathway. Genetics, 2021. 217(2):iyaa008.

39. Hsueh Y.P., Mahanti P., Schroeder F.C., and Sternberg P.W., Nematode-trapping fungi eavesdrop on nematode pheromones. Current Biology, 2013. 23(1):83–86.

40. Su H., Zhao Y., Zhou J., Feng H., Jiang D., Zhang K.Q. et al., Trapping devices of nematode-trapping fungi: formation, evolution, and genomic perspectives. Biological Reviews, 2017. 92(1):357–368.

41. Ma N., Zhao Y., Wang Y., Yang L., Li D., Yang J. et al., Functional analysis of seven regulators of G protein signaling (RGSs) in the nematode-trapping fungus Arthrobotrys oligospora. Virulence, 2021. 12(1):1825–1840.

42. Yang L., Li X., Ma Y., Zhang K., and Yang J., The Arf-GAP Proteins AoGcs1 and AoGts1 Regulate Mycelial Development, Endocytosis, and Pathogenicity in Arthrobotrys oligospora. Journal of Fungi, 2022. 8(5):463.

43. Van Ooij C., Hungry fungus eats nematode. Nature Reviews Microbiology, 2011. 9(11):766–767.

44. Schmid S.L., Reciprocal regulation of signaling and endocytosis: Implications for the evolving cancer cell. Journal of Cell Biology, 2017. 216(9):2623–2632.

45. Atanassov C.L., Muller C.D., Dumont S., Rebel G., Poindron P., and Seilers N., Effect of ammonia on endocytosis and cytokine production by immortalized human microglia and astroglia cells. Neurochemistry international, 1995. 27(4-5):417–424.

46. Abenza J.F., Pantazopoulou A., Rodríguez J.M., Galindo A., and Peñalva M.A., Long-distance movement of Aspergillus nidulans early endosomes on microtubule tracks. Traffic, 2009. 10(1):57–75.

47. Komatsu A., Iida I., Nasu Y., Ito G., Harada F., Kishikawa S. et al., Ammonia induces amyloidogenesis in astrocytes by promoting amyloid precursor protein translocation into the endoplasmic reticulum. Journal of Biological Chemistry, 2022. 298(5).

48. Schindler J.F., and de Vries U., Effects of ammonia, chloroquine, and monensin on the vacuolar apparatus of an absorptive epithelium. Cell and tissue research, 1990. 259(2):283–292.

49. Daussy C.F., and Wodrich H., “Repair Me if You Can”: membrane damage, response, and control from the viral perspective. Cells, 2020. 9(9):2042.

50. Blazek A.D., Paleo B.J., and Weisleder N., Plasma membrane repair: a central process for maintaining cellular homeostasis. Physiology, 2015. 30(6):438–448.

51. Wernet N., Wernet V., and Fischer R., The small-secreted cysteine-rich protein CyrA is a virulence factor participating in the attack of Caenorhabditis elegans by Duddingtonia flagrans. PLoS Pathog, 2021. 17(11):e1010028.

52. Jimenez A.J., Maiuri P., Lafaurie-Janvore J., Divoux S., Piel M., and Perez F., ESCRT machinery is required for plasma membrane repair. Science, 2014. 343(6174):1247136.

53. Gatta A.T., and Carlton J.G., The ESCRT-machinery: closing holes and expanding roles. Current opinion in cell biology, 2019. 59:121–132.

54. Suresh H.G., Pascoe N., and Andrews B., The structure and function of deubiquitinases: Lessons from budding yeast. Open biology, 2020. 10(10):200279.

55. Colot H.V., Park G., Turner G.E., Ringelberg C., Crew C.M., Litvinkova L. et al., A high-throughput gene knockout procedure for Neurospora reveals functions for multiple transcription factors. Proceedings of the National Academy of Sciences, 2006. 103(27):10352–10357.

56. Stiernagle T., Maintenance of C. elegans. 1999.

57. Yang J., Yu Y., Li J., Zhu W., Geng Z., Jiang D. et al., Characterization and functional analyses of the chitinase-encoding genes in the nematode-trapping fungus Arthrobotrys oligospora. Archives of microbiology, 2013. 195(7):453–462.

58. Livak K.J., and Schmittgen T.D., Analysis of relative gene expression data using real-time quantitative PCR and the 2^− ΔΔCT^method. methods, 2001. 25(4):402–408.

59. Ma Y., Yang X., Xie M., Zhang G., Yang L., Bai N. et al., The Arf-GAP AoGlo3 regulates conidiation, endocytosis, and pathogenicity in the nematode-trapping fungus Arthrobotrys oligospora. Fungal Genetics and Biology, 2020. 138:103352.

60. Liu X.H., Lu J.P., Zhang L., Dong B., Min H., and Lin F.C., Involvement of a Magnaporthe grisea Serine/Threonine kinase gene, Mg ATG1, in appressorium turgor and pathogenesis. Eukaryot Cell, 2007. 6(6):997–1005.

61. Wang C., Zhang S., Hou R., Zhao Z., Zheng Q., Xu Q. et al., Functional analysis of the kinome of the wheat scab fungus Fusarium graminearum. PLoS Pathog, 2011. 7(12):e1002460.

62. Tunlid A., Åhman J., and Oliver R., Transformation of the nematode-trapping fungus Arthrobotrys oligospora. FEMS microbiology letters, 1999. 173(1):111–116

